# Joint regulation of growth and division timing drives size homeostasis in proliferating animal cells

**DOI:** 10.1101/173070

**Authors:** Abhyudai Singh, Cesar A. Vargas-Garcia, Mikael Björklund

**Affiliations:** Department of Biomedical Engineering, University of Delaware, Newark, Delaware, USA; Department of Electrical and Computer Engineering, University of Delaware, Newark, Delaware, USA; Department of Mathematical Sciences,University of Delaware, Newark, Delaware, USA; Division of Cell and Developmental Biology, College of Life Sciences, University of Dundee, Dundee, UK.

## Abstract

How organisms maintain cell size homeostasis is a long-standing problem that remains unresolved, especially in multicellular organisms. Recent experiments in diverse animal cell types demonstrate that within a cell population the extent of growth and cellular proliferation (i.e., fitness) is low for small and large cells, but high at intermediate sizes. Here we use mathematical models to explore size-control strategies that drive such a non-monotonic fitness profile resulting in an optimal cell size. Our analysis reveals that if cell size grows exponentially or linearly over time, then fitness always varies monotonically with size irrespective of how timing of division is regulated. Furthermore, if the cell divides upon attaining a critical size (as in the Sizer or size-checkpoint model), then fitness always increases with size irrespective of how growth rate is regulated. These results show that while several size control models can maintain cell size homeostasis, they fail to predict the optimal cell size, and hence unable to explain why cells prefer a certain size. Interestingly, fitness maximization at an optimal size requires two key ingredients: 1) The growth rate decreases with increasing size for large enough cells; and 2) The cell size at the time of division is a function of the newborn size. The latter condition is consistent with the Adder paradigm for division control (division is triggered upon adding a fixed size from birth), or a Sizer-Adder combination. Consistent with theory, Jurkat T cell growth rates, as measured via oxygen consumption or mitochondrial activity, increase with size for small cells, but decrease with size for large cells. In summary, regulation of both growth and cell division timing is critical for size control in animal cells, and this joint-regulation leads to an optimal size where cellular fitness is maximized.

Address inquires to A. Singh.

---

Size control is a fundamental aspect of biology that is seen at every level of organization, but in most cases remains poorly understood [1–5]. One simple solution for size control is the Sizer or size-checkpoint model that couples cell-cycle transitions to attainment of a minimum cell mass or size. An alternative solution is the Adder model, where cells add a fixed amount of size in each division cycle independent of daughter size. While strong evidence for an Adder has been reported in diverse prokaryotes [6–16], and budding yeast [17], it remains to be seen if similar size homeostasis mechanisms are at play in higher eukaryotes.

Recent experiments in proliferating animal cells reveal an intriguing observation about cellular fitness that ties deeply into size control as this suggested why cells aim to maintain certain size. In [18], fitness was assayed by first sorting a cell population into several subpopulations based on size, and then measuring the net proliferation (increase in cell count) for each cultured subpopulation after 72 hrs. Interestingly, data across several cell types shows a non-monotonic bell-shaped fitness profile, where fitness is maximized at an optimal cell size (Fig. 1). Importantly, apoptosis rates were similar for large and average-sized cells, and hence the decrease in fitness at higher sizes is not due to elevated cell death [18, 19]. Finally, cellular proliferation measured via dye-dilution experiments shows identical trends for varying cells sizes (see supplementary figure S4 in [18]), providing an independent confirmation for the existence of an optimal cell size where proliferative capacity is maximal. While the data in Fig. 1 is collected on asynchronous cell populations, one could consider a modified fitness vs. size assay that is restricted to newborn cells. Since large newborns will have lower proliferation compared to randomly chosen large cells that are further along the cell cycle, the dip in fitness at higher sizes will be even more pronounced for newborns. In contrast, randomly chosen small cells are presumably at the beginning of their cell cycle, and have similar proliferation as compared to small newborns. Thus, measuring cellular proliferation with varying newborn size not only preserves the bell-shaped fitness profile, but it also leads to a clear mathematical formulation of fitness. More specifically, let the function *T*(*V_b_*) denote the average cell-cycle duration for a cell born with size *V_b_*. Then, fitness is simply defined by

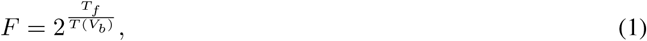

where *T_f_* is the fixed duration of the experiment. While cell-cycle lengths are generally modulated to decrease with newborn size [21, 22], a bell-shaped fitness curve requires *T*(*V_b_*) to decrease with increasing *V_b_* for small cells, but increase with *V_b_* for large cells. Our analysis below shows that most models for size homeostasis cannot capture this non-monotonic behavior, and we identify selected scenarios that are consistent with it. Having setup the problem, we introduce mathematical models for cell size control and investigate *F* vs. *V_b_* profiles.

**Figure 1:**
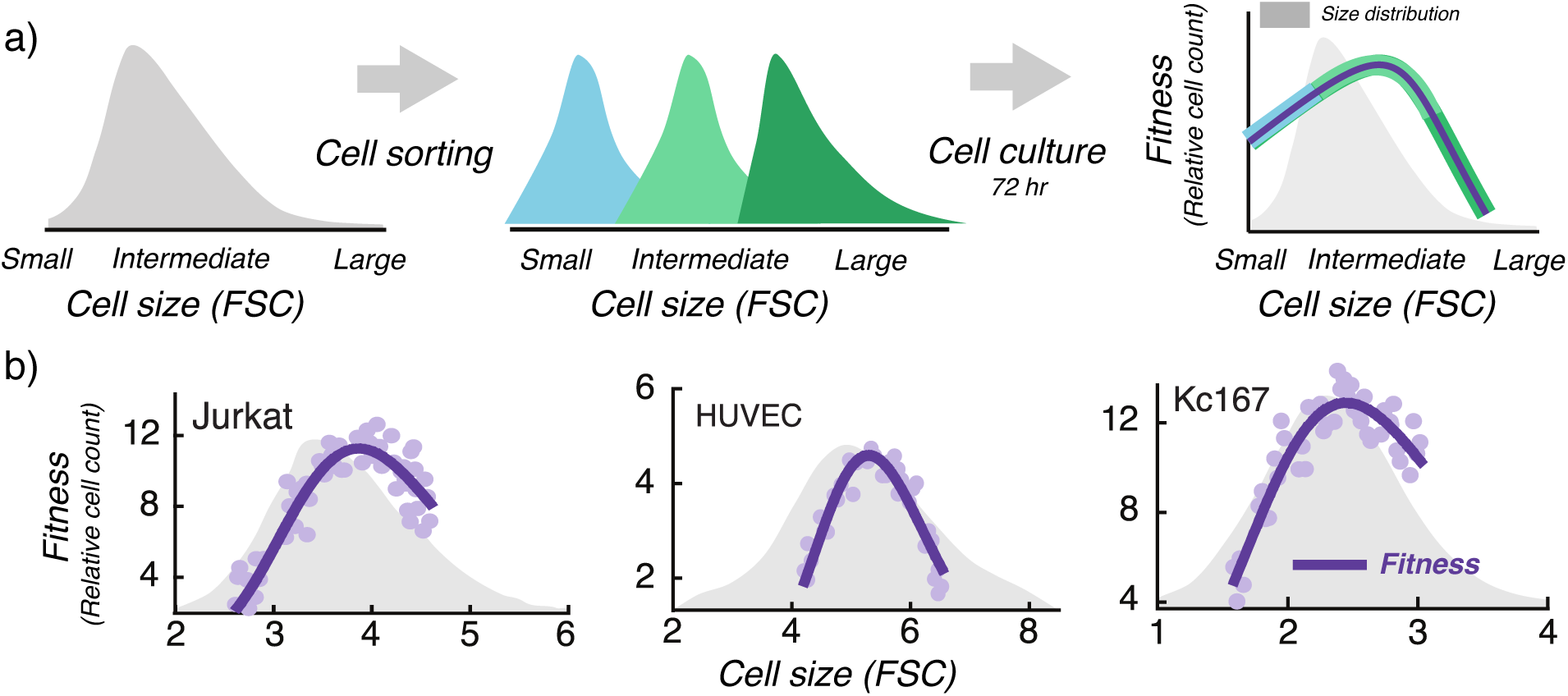
Cellular fitness is maximized at an optimal cell size. a) Using forward scatter intensity (FSC) as a proxy for cell size [18, 20], flow cytometry is used to sort an original unsynchronized cell population (grey) into several subpopulations with different cell sizes. Each subpopulation is cultured for 72 hrs (approximately 3 – 5 cell generations), and fitness is quantified by measuring the relative change in cell count. Interested readers are referred to the material and methods of [18] for further details. b) Measured fitness is plotted as a function of the average subpopulation FSC at the time of sorting for three different cell types: Jurkat cells (human T lymphocyte cell line), HUVEC (human umbilical vein endothelial cells; a primary cell line) and Kc167 (a widely used Drosophila cell line). Original cell size distribution is shown in grey.

The dynamics of cell size is described by

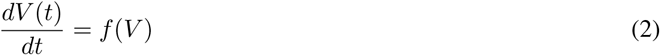

where, *V*(*t*) is the size of an individual cell at time *t* since the start of cell cycle, and the function *f* describes a general size-dependent growth rate. As in prior work, regulation of the cell-cycle time *T* is conveniently described by a relationship between the newborn size *V_b_* := *V*(0), and the size just prior to mitosis *V_d_* := *V*(*T*),

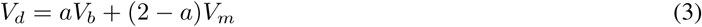

with parameters *V_m_* and *a* ∈ [0, 2] [17, 23–25]. Assuming division of a mother cell into two daughters, the upper bound *a* ≤ 2 is needed for system stability (boundedness of newborn sizes over successive generations), while the lower bound 0 ≤ *a* ensures positivity of *V_d_*. Specific values of *a* correspond to well known division control strategies:

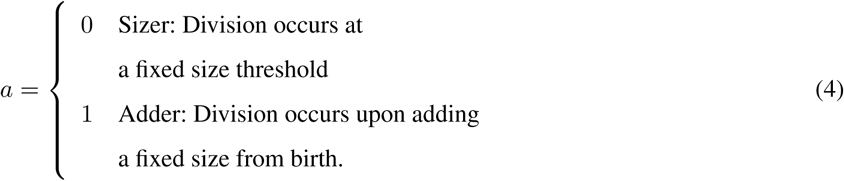

Values of *a* between zero and one imply an adder-sizer mixture, and such combinatorial control of cell size have been proposed in many organisms [24, 26–28]. *V_m_* is the mean newborn size, and (3) ensures size doubling from birth to division for an average newborn, i.e., *V_d_* = 2*V_m_* when *V_b_* = *V_m_*. Finally, we point out that exponential growth in cell size (*f* (*V*) ∝ *V*) coupled with a timer, i.e., division occurs after a fixed time from cell birth, corresponds to *a* = 2. However, this simplistic case is non-homeostatic, in the sense that, the variance in newborn size grows unboundedly over time in the presence of noise [29, 30].

The Sizer, where mitosis is triggered upon reaching a prescribed size threshold, is perhaps the simplest (and the oldest proposed) mechanism for size homeostasis. In this case, the cell-cycle time

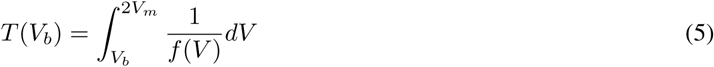

monotonically decreases with increasing *V_b_* for any function *f*. As a consequence, the proliferative fitness *F* is a monotonically increasing function of *V_b_*. Thus, Sizer-based division (*a* = 0) is inconsistent with non-monotonic cellular proliferation (Fig. 1), irrespective of how cellular growth is regulated [31]. As a corollary to this result, a strictly positive *a* that leads to *V_d_* being dependent on *V_b_* is a necessary determinant of an optimal cell size. Given the constraints on the values *a* can take, we next explore if similar constraints arise on the growth rate.

Assume exponential growth

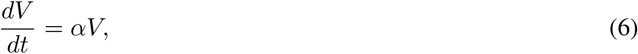

where *α* is the exponential growth coefficient. In this case,

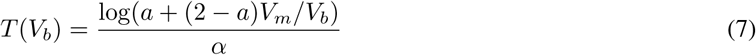

monotonically decreases with *V_b_* for all values of *a* ∈ [0,2], i.e., *F* increases with size irrespective of how division timing is regulated (Fig. 2a). An important implication of this result is that while exponential growth coupled with some form of division control may explain size homeostasis in prokaryotes and microbial eukaryotes, it does not yield the size optimality seen in animal cells (Fig. 1). Now consider a generalization of exponential growth to any monotonically increasing growth rate *f* (*V*). After a change in variable *V* = *aV_b_* + (2 – *a*)*v*, the cell-cycle time can be rewritten as

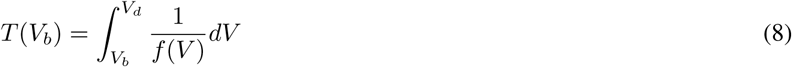

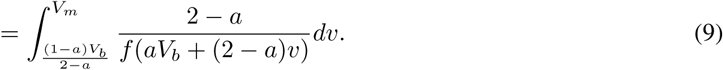

**Figure 2:**
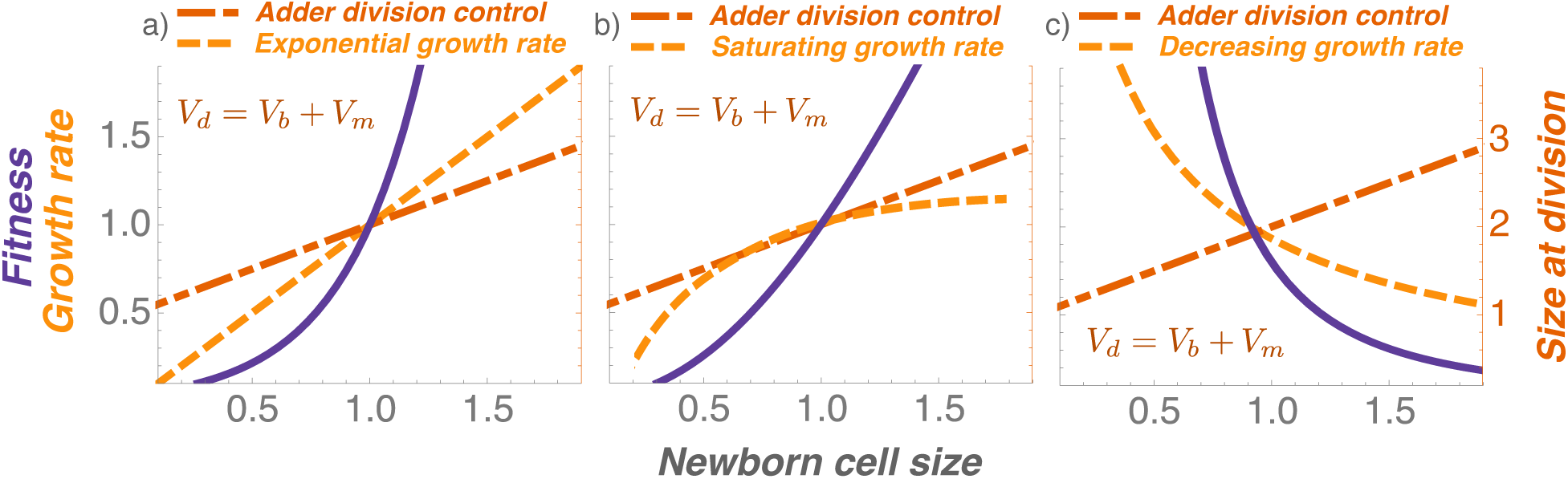
Monotonic growth rates coupled with Adder-based division control yield monotonic fitness profiles. Cell size at division as per (3), the growth rate function in (2), and the corresponding fitness profile computed via (1) & (8) is plotted for different size homeostasis mechanisms. Exponential growth (a), or an increasing growth rate followed by saturation (b) together with the Adder model always lead to cellular fitness monotonically increasing with size. In contrast, a decreasing growth rate together with an Adder yields a decreasing fitness profile (c). Newborn cell size (x axis) is normalized by its mean. Fitness, growth rate, and size at division (y axes) are normalized by their corresponding values for the mean newborn size.

For 0 ≤ *a* ≤ 1, the lower-limit of the definite integral is an increasing function of *V_b_*, while the function being integrated is a decreasing function of *V_b_*. Thus, for any arbitrary increasing function *f*, *T* decreases with newborn size for any Sizer-Adder combination. Interestingly, non-monotonicity in *T* can arise for values of *a* much larger than one. We illustrate this point by a saturating growth function that has been reported for mouse lymphoblast cells [32]

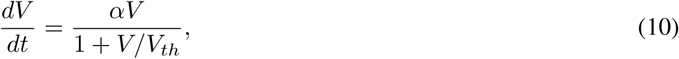

with an additional parameter *V_th_* > 0. While fitness monotonically increases with newborn size for *a* ≤ 1 as predicted above (Fig. 2b), it becomes non-monotonic for *a* > 1 (Fig. 3a). Intuitively, for large cells (*V* ≫ *V_th_*), growth is linear and

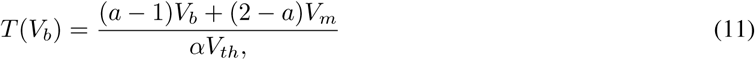

**Figure 3:**
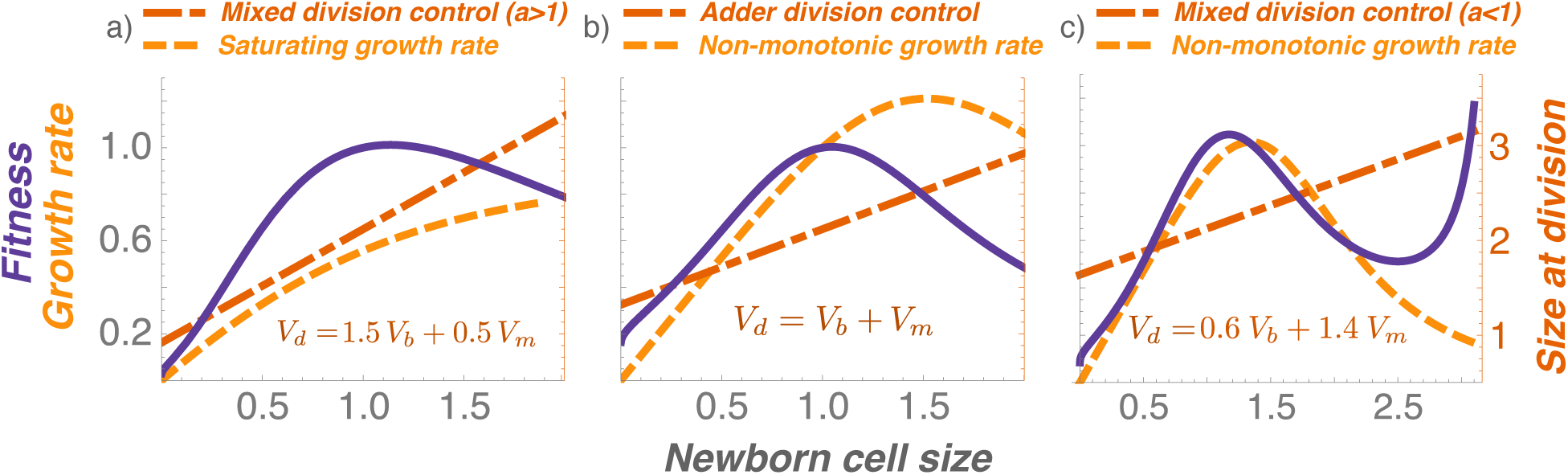
A non-monotonic growth rate coupled with Adder-based division control maximizes fitness at an optimal cell size. a) A monotonically increasing growth rate can drive a bell-shaped fitness profile if *a* > 1, i.e., size added in a cell-cycle duration increases with daughter cell size. b) A non-monotonic growth rate together with an Adder (*a* = 1) yields an optimal cell size consistent with data in Fig. 1. c) A non-monotonic growth rate together with an Adder-Sizer combination (*a* < 1) results in a complex profile: a bell-shaped fitness for most physiological cell sizes, but fitness again increases at higher sizes. Newborn cell size (x axis) is normalized by its mean. Fitness, growth rate, and size at division (y axes) are normalized by their corresponding values for the mean newborn size.

become longer (and hence, decreasing proliferation) with increasing newborn size iff *a* > 1. It is interesting to note that a decreasing growth rate with size can lead to contrasting results for a Sizer and Adder - while as per (5), *F* is an increasing function of *V_b_* for a Sizer, it becomes a decreasing function for an Adder (Fig. 2c). This result motivate a non-monotonic growth rate

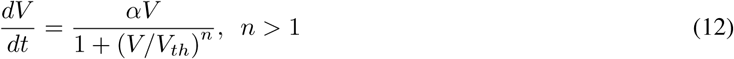

where growth is exponential for small newborns and *F* increases with size. However, for large newborns, growth rate is decreasing (∝ *V*^1–*n*^), and *F* decreases with size if division timing is appropriately controlled. Our analysis reveals that *a* ≥ 1 (that includes the adder mechanism) combined with (12) guarantees a non-monotonic fitness curve (Fig. 3b). Interestingly, the non-monotonicity is preserved for values of *a* smaller than one, but with an additional feature - fitness again increases for very large newborns (Fig. 3c). While such an increase is not seen in the data, it is possible that sizes needed for this effect fall outside the physiological range.

Overall, our results suggest size-based regulation of *both* growth and timing of division as necessary determinants of an optimal cell size. This joint regulation is not sufficient, as any monotonically increasing growth rate, coupled with any Sizer-Adder combination leads to cellular fitness increasing with size. Mathematical models identify two scenarios for a bell-shape fitness profile. The first scenario involves a saturating growth function (exponential growth for small cells, linear growth for large cells). While such a growth rate together with a Timer (size-independent cell-cycle length) is sufficient for size homeostasis [29, 30], existence of an optimal cell size requires division timing to be controlled such that *a* > 1 in (3), i.e., size added in each division cycle increases with newborn size (Fig. 3a).

The second scenario considers a non-monotonic growth rate (12) that decreases with size for sufficiently large cells, as has been reported for some mammalian cells [18, 33–35]. This lowering of growth rate in larger cells can be attributed to a decreased surface area-to-volume ratio leading to insufficient nutrient exchange for supporting growth [36, 37]. Indeed, Jurkat T cell growth rates, as measured via oxygen consumption and mitochondrial activity, increase with size for small cells, but decrease with size for large cells [18]. Note that non-monotonic growth combined with a Sizer or Timer will lead to an increasing or flat fitness vs. size profile, respectively. However, (12) in conjunction with an Adder (Fig. 3b) or a Sizer-Adder mixture (Fig. 3c) is sufficient to drive fitness optimality at intermediate sizes. Consistent with this result, recent single-cell tracking of several mammalian cell types shows size added in a cell-cycle duration to be independent of the newborn size as per the Adder model [38]. In summary, our results uncover novel insights into size control principles, and provide a mechanistic explanation for the existence of an optimal size in proliferating animal cells.

## Acknowledgments

This work is supported by a grant from the National Institute of Health to AS (No. 1R01GM126557-01).

